# *In vivo* optochemical control of cell contractility at single cell resolution by Ca^2+^-mediated light-activation of myosin (CaLM)

**DOI:** 10.1101/255372

**Authors:** Deqing Kong, Zhiyi Lv, Matthias Häring, Fred Wolf, Jörg Großhans

## Abstract

The spatial and temporal dynamics of cell contractility plays a key role in tissue morphogenesis, wound healing and cancer invasion. Here we report a simple, single cell resolution, optochemical method to induce minute-scale cell contractions *in vivo* during morphogenesis. We employed the photolabile Ca^2+^ chelator *o*-nitrophenyl EGTA to induce bursts of intracellular free Ca^2+^ by laser photolysis. Ca^2+^ bursts appear within seconds and are restricted to individual target cells. Cell contraction reliably followed within a minute, to about half of the cross-sectional area. Increased Ca^2+^ levels and contraction were reversible and the target cells further participated in tissue morphogenesis. Depending on Rho kinase (Rok) activity but not RhoGEF2, cell contractions are paralleled with non-muscle myosin-II accumulation in the apico-medial cortex, indicating that Ca^2+^ bursts trigger non-muscle myosin II activation. Our approach can be easily adapted to many experimental systems and species, as no specific genetic elements are required and a widely used reagent is employed.

## Introduction

Contractility underlies manifold processes in cell and tissue morphogenesis, including cell migration, cell shape changes, or junction collapse [1-4]. In epithelial tissues, cell contractions impact neighboring cells by exerting forces on adherens junctions. This mechanical linkage may elicit specific responses and could thus positively or negatively affect contractility and cytoskeletal organization in neighboring cells i.e. mediate non-autonomous mechanical behaviors [5]. Within a tissue, cellular contraction and cell-cell interactions based on such force transduction can contribute to emergent tissue behavior, such as the formation of folds and furrows. The function of mutual cell-cell interactions, however, is difficult to study by classical genetic approaches. What is needed are methods for acute noninvasive interventions with high temporal and spatial resolution, ideally on the scale of seconds and of single cells.

For controlling cell contractility optogenetic approaches have recently been developed. Cell contractility can be inhibited by optically induced membrane recruitment of PI(4,5)P_2_ leading to interference with phosphoinositol metabolism and subsequent suppression of cortical actin polymerization [6]. Optical activation of contractility has been achieved by light-induced activation of the Rho-Rok pathway, which controls Myosin II based contractility [7,8]. While functionally effective, such optogenetic methods require multiple transgenes driving the expression of modified proteins such as light-sensitive dimerization domains, which restricts the application to genetically tractable organisms. In addition, chromophores used in optogenetic effectors are activated by light in the visible spectrum, which limits the choice of labels and reporters for concurrent cell imaging.

Optochemical methods represent an alternative to genetically encoded sensor and effector proteins [9]. Intracellular calcium ions (Ca^2+^) are known to be an important regulator of contractility in many cell types. Ca^2+^ not only plays a central role in muscle contraction, but also in cultured epithelial cells [10], in amnioserosa cells during dorsal closure [11], during neural tube closure [12,13] and in the folding morphogenesis of the neural plate [14]. In *Drosophila* oogenesis, tissue wide increase of intracellular Ca^2+^ activates Myosin II and impairs egg chamber elongation [15]. In *Xenopus,* a transient increase in Ca^2+^ concentration induces apical constriction in cells of the neural tube [16]. Although the detailed mechanism of Ca^2+^ induced contraction in non-muscle cells remains to be resolved, it conceivably offers a simple and temporally precise way to interfere with and control contractile activity. In neuroscience, optochemical methods for the release of intracellular Ca^2+^ have been well established and widely employed [17]. Here we report a Ca^2+^ uncaging method to control epithelial cell contractility on the scale of seconds and at single cell resolution during tissue morphogenesis in *Drosophila* embryos. Optochemical control of contractility by Ca^2+^ uncaging has minimal spectral overlap with fluorescent protein reporters and optogenetic activators. Our results provide evidence for a Rok dependent effect of increased intracellular Ca^2+^ on activating non-muscle myosin II and its recruitment to the actomyosin cortex.

## Results

### Uncaging induces a rapid Ca^2+^ burst in epithelial cells in *Drosophila* embryos

Photolysis of the Ca^2+^ chelator *o*-nitrophenyl EGTA (NP-EGTA) [18] (Fig. 1A) is widely used in neurobiology for the modulation of intracellular Ca^2+^ concentration [19]. Here, we employed the membrane-permeant acetoxymethyl (AM) ester derivative, which complexes Ca^2+^ once the AM moiety is cleaved off by intracellular esterases. Following microinjection into staged embryos, uncaging was induced in the focal volume of a pulsed 355 nm laser beam (Fig. 1B). To allow for concomitant uncaging and imaging, we used a setup, in which the light paths of the UV laser for uncaging and the excitation laser for confocal imaging in the visible spectrum were controlled independently. We conducted experiments in the lateral epidermis of *Drosophila* embryos during germband extension (stage 7). The epidermis during this stage constitutes a columnar epithelium with a cell diameter in the range of about 8 μm and cell height of about 25 μm (Fig. 2A).

**Figure 1.**
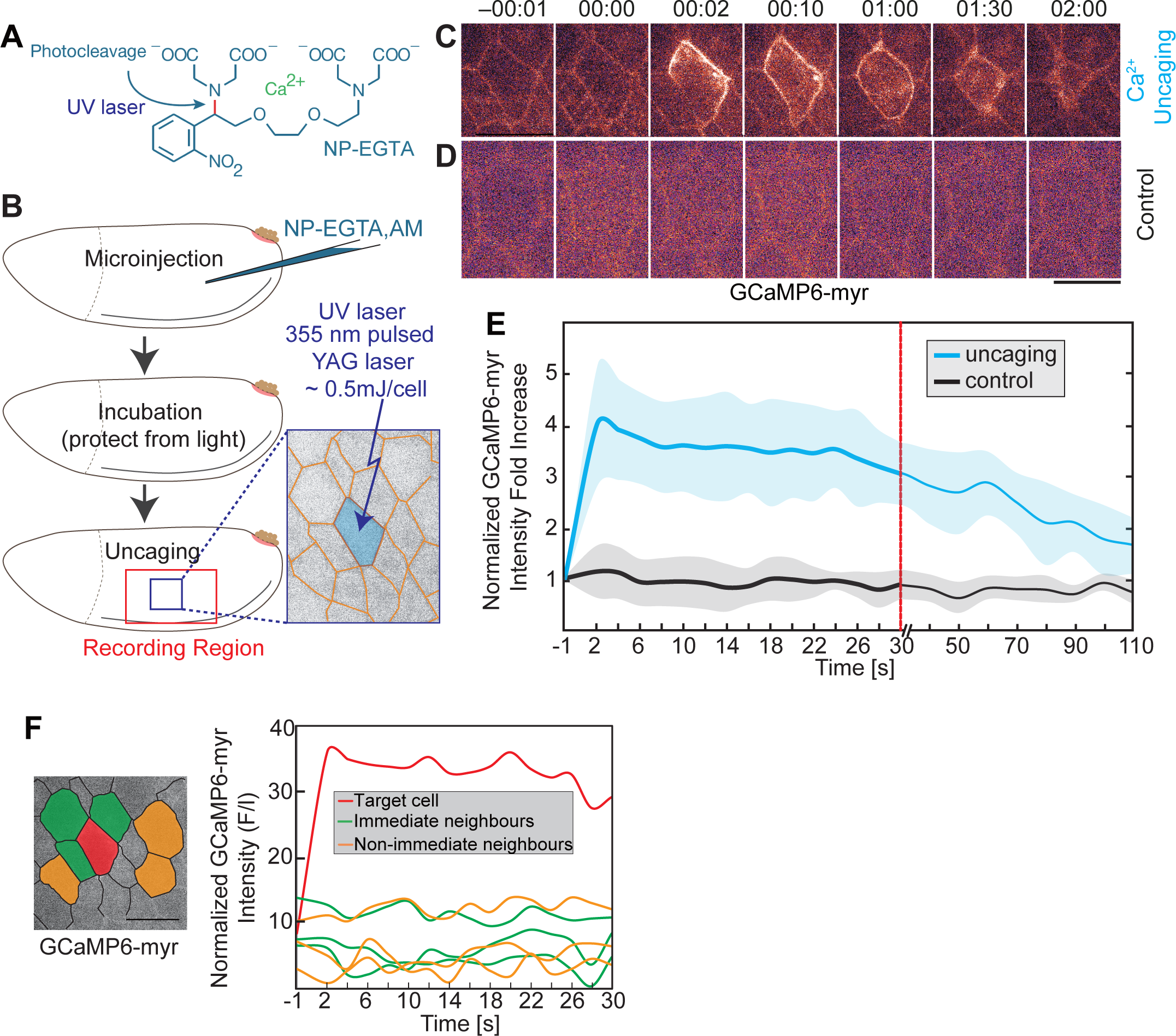
**Uncaging induces rapid increase in intracellular Ca**^**2+**^ **concentration in epithelial target cells.** **(A)** Structure of the cage compound NP-EGTA. UV illumination cleaves the compound and releases free Ca^2+^. The photo-labile bond is indicated in red. **(B)** Experimental scheme for Ca^2+^ uncaging in *Drosophila* embryos. NP-EGTA,AM was injected into the staged embryos. Followed by a short incubation, a selected cell was exposed to a UV laser flash. Target cell is indicated in blue. **(C, D)** Images from time lapse recording of embryos (stage 7, lateral epidermis) expressing a membrane bound Ca^2+^ sensor (GCaMP6-myr) and injected with (C) 2 mM NP-EGTA,AM or (D) with buffer (control). Time in minutes:seconds **(E)** Normalized fluorescence intensity of GCaMP-myr in the target cell. Mean (bold line, 3 cells in 3 embryos) with standard deviation of the mean (ribbon band). **(F)** Normalized fluorescence intensity of GCaMP sensor in target cell (red), 3 next neighbors (green) and 3 non-immediate neighbors (orange). Scale bars 10 μm.

**Figure 2.**
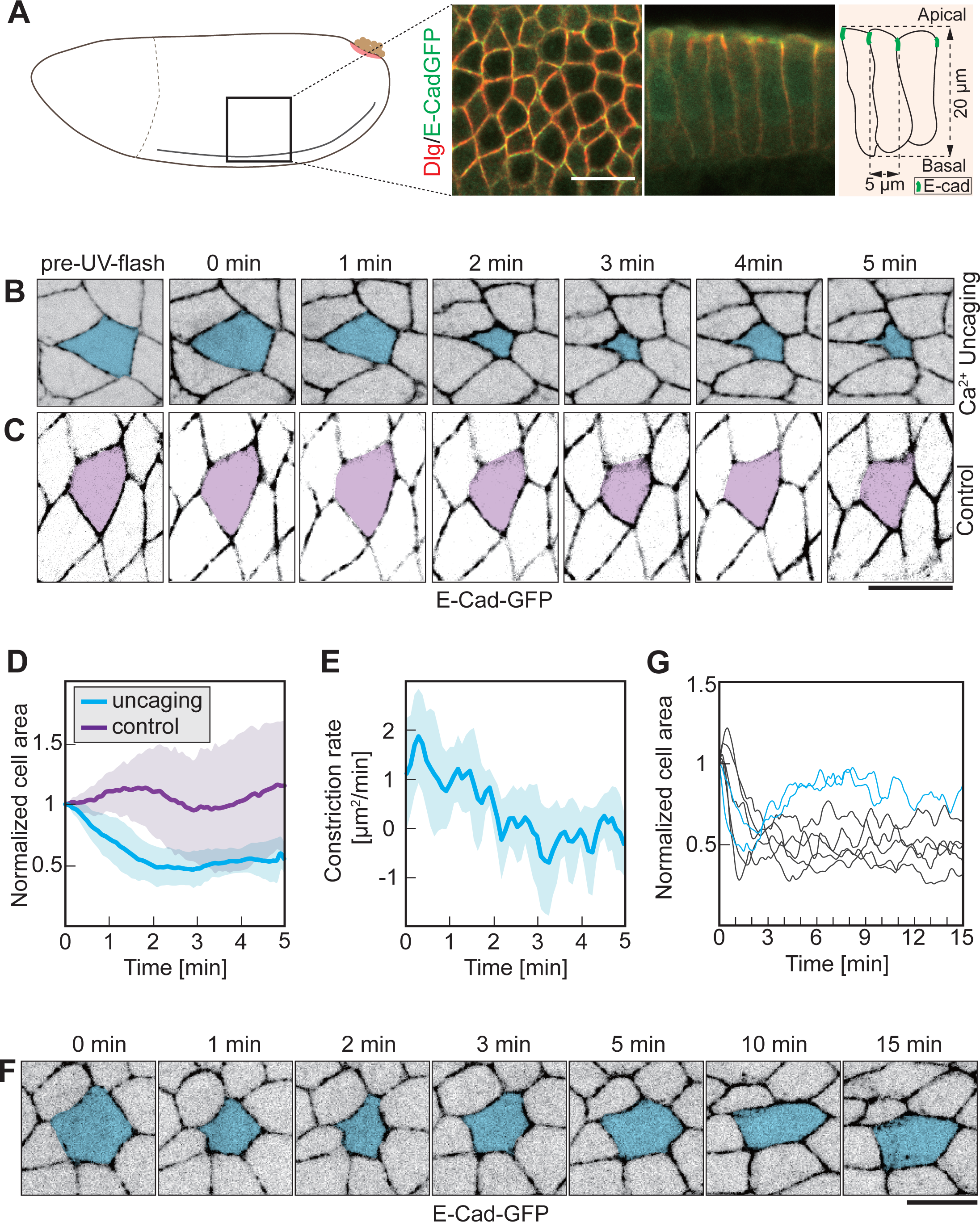
**Ca**^**2+**^ **uncaging triggers apical-constriction in a columnar epithelium.** **(A)** Schematic drawing and morphology of columnar epithelium in the lateral epidermis in stage 7 *Drosophila* embryos. **(B, C)** Images from a time lapse recording embryos expressing E-Cad-GFP and injected with (B) 2 mM NP-EGTA,AM or (C) buffer and exposed to the UV laser. Target cells are labelled in blue or purple. **(D)** Cross-sectional area of target cells over time. Cell areas were normalized by the initial size (the first frame of recording after uncaging). Mean (bold line) with standard deviation of the mean (ribbon band). Uncaging, (blue) 8 cells in 8 embryos. Control (purple) 5 cells in 5 embryos. **(E)** Apical constriction rate over time for the Ca^2+^-uncaging-activated contracting cells in D (n= 8). Mean (bold line) with standard deviation of the mean (ribbon bands). **(F)** Images from time lapse recording showing long term behavior after uncaging. Target cell is marked in blue. **(G)** Cross-sectional area of target cells over 15 minutes after Ca^2+^ uncaging. Cell contraction in 2 out of 7 target cells was reversible (blue lines). Scale bars 10 μm.

We recorded changes of intracellular Ca^2+^ concentration induced by uncaging using a genetically encoded Ca^2+^ sensor protein, GCaMP6s. Embryos expressing a membrane bound, myristoylated variant of GCaMP6s [20] were injected with NP-EGTA-AM and subjected to uncaging. We observed a transient increase of GCaMP6 fluorescence within a second specifically in cells targeted by a UV light pulse (Fig. 1C Supp. movie 1). Quantification of GCaMP fluorescence (ΔF/F0) showed a fourfold increase within two seconds. Afterwards, GCaMP6s fluorescence gradually decreased to near initial levels within a few minutes (Fig. 1E). As GCaMP6s has a decay time constant in the range of seconds, this indicates that Ca^2+^ clearance and extrusion mechanisms in the epithelial cells operate on an effective time scale of minutes. We did not detect an increase of GCaMP6s fluorescence after UV exposure in control embryos injected with buffer only (Fig. 1D E).

The increase of the Ca^2+^ sensor signal was restricted to the individual target cell (Fig. 1C Supp. movie 1). The Ca^2+^ sensor signal in the next and next-next neighbors of the target cell was temporally constant and comparable to control embryos (Fig. 1F). In summary, our experiments show that Ca^2+^ uncaging with single cell precision can be conducted in epithelial tissue in *Drosophila* embryos. Uncaging leads to a reversible, second-scale increase of intracellular Ca^2+^ concentration that is restored by cell intrinsic mechanisms on a minute scale. The magnitude of the Ca^2+^ increase was similar to what was previously reported for neuronal cells [21].

### Ca^2+^ bursts induce cell contraction

We next investigated the consequence of Ca^2+^ bursts on cell shape. We conducted uncaging in embryos expressing E-Cad-GFP, which labels adherens junctions near the apical surface of the epithelium (Fig. 2A). We detected a contraction of the target cell to about half of the apical cross-sectional area following uncaging (Fig. 2B and S1A, Supp. movie 2 and 3). Target cells in control embryos injected with buffer remained largely unaffected (Fig. 2C). Quantification revealed a reduction by half of the cross sectional area within 1–2 min in the target cell but not in controls (Fig. 2C and S1). Most cells remained contracted during the following 15 min, whereas a minority of cells reexpanded to the original cross-sectional area (Fig. 2F, G). We did not observe that the exposure to UV laser and Ca^2+^ uncaging noticeably affected the further behaviour of the target cells and surrounding tissue. Cells showed typical oscillations with periods of a few minutes and amplitudes of 10–20% (Fig. 2F, G) [22,23]. We did not observe that target cells were extruded or got lost from epithelia tissue. This behavior indicates that the Ca^2+^ uncaging is compatible with ongoing tissue morphogenesis.

### Induced cell contraction in a squamos epithelium

Next we applied Ca^2+^ uncaging to a different tissue in *Drosophila* embryos. The amnioserosa represents a squamos epithelium on the dorsal side of the embryo with cells about 15 μm im diameter and only 3 μm in height (Fig. 3A–C). As in the lateral epidermis we employed E-Cadherin-GFP to label the apical cell outlines (Fig. 3B). Ca^2+^ uncaging led to contraction of the target cell but not its neighbors (Fig. 3D Supp. movie 4). Quantification of the apical cross sectional areas revealed specific uncaging induced contraction within a minute (Fig. 3E). The peak constriction rate was observed about 30 s after uncaging (Fig. 3F). In summary, our experiments show that Ca^2+^ uncaging can be employed as a noninvasive method to induce contractions in selected single cells in different cell types and tissues.

**Figure 3.**
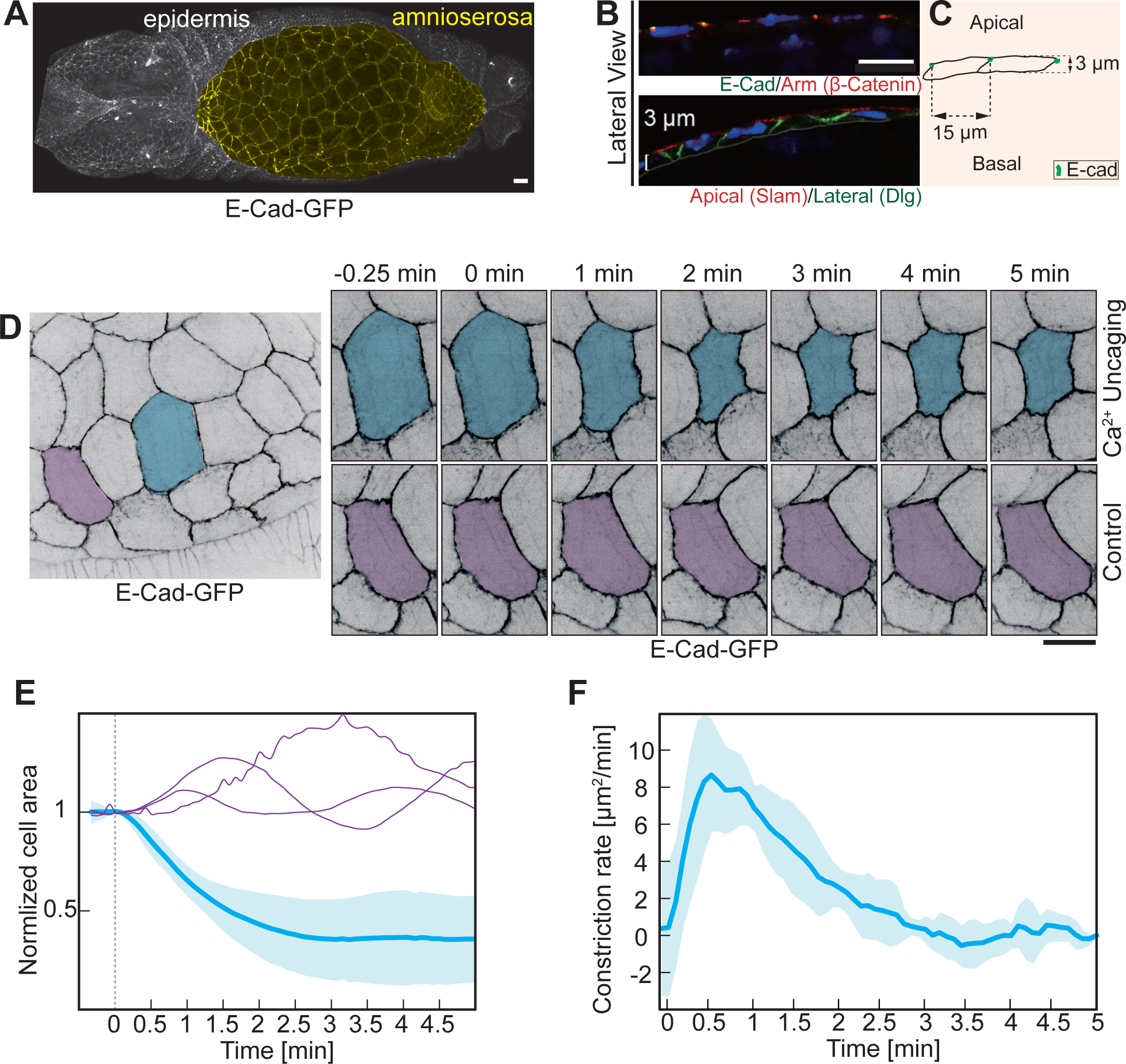
**Ca**^**2+**^ **uncaging triggers apical-constriction in a squamous epithelium.** **(A–C)** Amnioserosa represents a squamous epithelium. Confocal image of Drosophila embryo expressing E-Cadherin-GFP. Amnioserosa labelled in yellow (A). Sagittal sections of amnioserosa cells. Confocal images (B) and schematic drawing (C). **(D)** Images from a time lapse recording in embryos (stage 14) expressing E-Cad-GFP and injected with 1 mM NP-EGTA,AM. target cell marked in blue. The control cell (next-next neighbor) marked in purple was not exposed to UV light. **(E)** Cross-sectional area of target cells and control cells normalized to initial size (the first frame of recording after uncaging). Mean (bold line) with standard deviation of the mean (ribbon band) (n=4 cells in 4 embryos). Cross-sectional area traces of 3 individual control cells are indicated in purple. **(F)** Apical constriction rate in target cells (n=4). Mean (bold line) with standard deviation of the mean (ribbon band). Scale bars 10 μm.

### Role of Myosin II in Ca^2+^ induced cell contraction

Multiple mechanisms are conceivable for Ca^2+^ induced cell contraction. Given their time scale in the minute range, it is unlikely that slow transcriptional or translational processes are involved. It is also unlikely that Ca^2+^ directly activates contraction similar to its role in muscle cells due to the distinct organization of cortical actomyosin and indicated by the substantial time lag between Ca^2+^ increase and cell contraction. Conceivably, the reduction in cell cross sectional area may involve Ca^2+^ induced osmotic changes and associated efflux of water, for example. Alternatively, Ca^2+^ may activate myosin II, similar to what has been reported for the *Drosophila* egg chamber[15]. Such a specific Myosin II activation may be mediated via Rho-Rok signaling or via Ca^2+^ dependent protein kinases or phosphatases, such as myosin light chain kinase (MLK) [24].

As a first step towards identifying the mechanism of Ca^2+^ induced cell contraction, we imaged myosin II dynamics following uncaging in embryos expressing E-Cad-GFP to label cell-cell contacts and sqh-mCherry (myosin regulatory light chain). sqh-mCherry fluorescence is a direct indicator of active myosin II mini filaments, which are visible as clusters. Myosin II is found associated with adherens junctions (junctional pool) and at the apical cortex (medial pool), where it is responsible for apical constriction [25]. We focused on the medial pool of myosin II. We observed an increase of sqh-mCherry fluorescence after about 0.5–1 min specifically in target cells (Fig. 4A–C). To establish a link between the increase in myosin II and the reduced cell area, we correlated both parameters with each other (Fig. 4D, E). Indeed, we detect a strong correlation, that the cell area is the smaller the higher the Myosin II activity.

**Figure 4.**
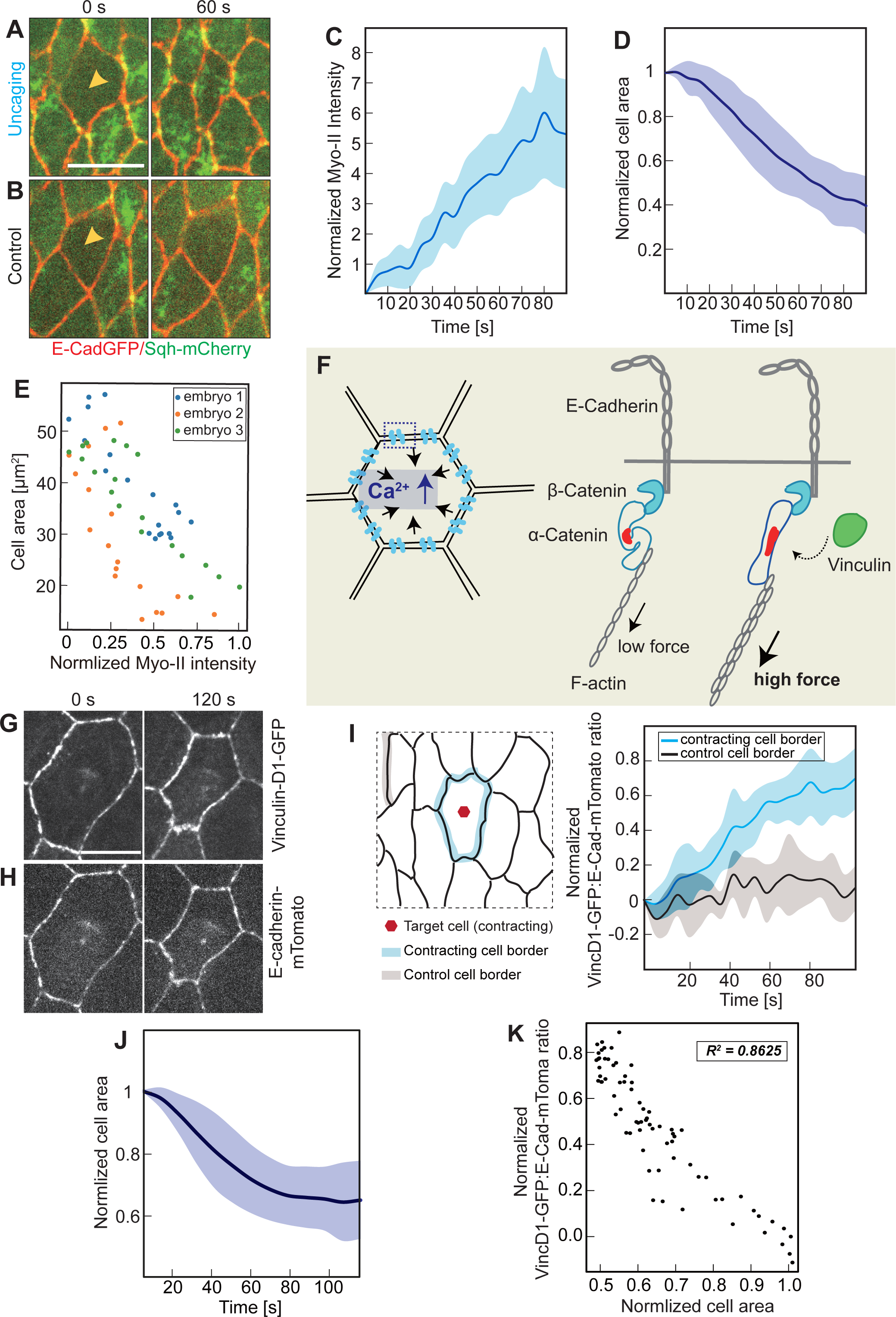
**Induction of myosin II and cortical tension by Ca**^**2+**^ **uncaging.** **(A–E)** Myosin II activation in target cells. **(A, B)** Embryos expressing Sqh-mCherry (green) and E-Cadherin-GFP (red) were injected with 2 mM NP-EGTA,AM (A) or buffer (B). Images from a time-lapse recording in cells of the lateral epidermis (stage 7). Time after UV illumination. Yellow arrowheads point to target cells. **(C)** medio-apical Sqh-mCherry fluorescence in target cells normalized to the fluorescence intensities in control cells. Mean (bold line) with standard deviation of the mean (ribbon band) (n=3 cells in 3 embryos). **(D)** Cross sectional area of target cells normalized to the initial area (the first frame of recording after uncaging). Mean (bold line) with standard deviation of the mean (ribbon band) (n=3 cells in 3 embryos). **(E)** Scatter plot of normalized medio-apical myosin II with cross-section area in target cells. **(F-K)** Forces increase in Ca^2+^-uncaging-activated constricting cells. **(F)** Schematic drawing of force dependent Vinculin association to adherens junctions and principle of the Vinculin reporter. **(G, H)** Images from time lapse recording of an amnioserosa cell after Ca^2+^ uncaging in embryos (stage 14) expressing E-Cad-mTomato and VinculinD1-GFP. **(I)** Ratio of VinculinD1-GFP and E-Cadherin-mTomato fluorescence the the junctions of the target and control cell (n=3 constricting cells and 9 inactive cell borders). **(J)** Cross-sectional area in target cells normalized to initial size (the first frame of recording after uncaging). Mean (bold line) with standard deviation of the mean (ribbon band) (n=3 cells in 3 embryos). **(K)** Scatter plot of normalized area of target cells with the VinculinD1/E-cadherin ratio. Scale bars 10 μm.

One expects that a contracting cell applies a force on the junctional complexes linking it to its neighbors within the epithelium (Fig. 4F). To assess this action, we employed a reporter for tension across adherens junctions, based on the force-dependent conformational state of α-Catenin [26-29]. α-Catenin exhibits a force-dependent switch between two stable conformations. In the closed state α-Catenin is bound to the Cadherin complex but does not bind to the D1 domain of Vinculin, because the central mechanosensitive modulatory (M) domain is inaccessible. In contrast, the central mechanosensitive modulatory (M) domain is exposed, when a force is applied to the molecule. α-Catenin bridges the Cadherin complex with the actin cytoskeleton and can thus sense and transduce forces acting on the adherens junctions. We thus introduced a GFP reporter based on the D1 domain of Vinculin together with E-Cadherin-tomato inserted at the endogenous locus (Fig. 4G and 4H, Supp. movie 5). We quantified the dynamics of VinD1-GFP fluorescence during an uncaging experiment (Fig. S4). We detected a significant increase in the range of 10% of reporter fluorescence at the junctions next to the contracting target cell in the time scale of a minute. We did not detect such an increase at distant junctions, which served as a control in this experiment. As the time scale in response to uncaging by area change and VincD1 reporter fluorescence were comparable we quantified their relationship (Fig. S4). The Vinc/E-cad ratio has been reported to correlate with junctional tension in *Drosophila* embryos [30]. We therefore quantified the dynamics of VincD1/E-cad fluorescence ratio in the Ca^2+^-uncaging-activated contracting cells (Fig. 4I–K). We detect an obvious increase of VincD1/E-cad fluorescence ratio at the junctions next to the contracting target cell that appeared on a time minute scale. We did not detect such an increase at distant junctions, which served as a control in this experiment. As the time scale in response to uncaging by area change and VincD1/E-cad fluorescence ratio were comparable we quantified their relationship. We plotted the Vinc/E-cad ratio against cell area from 3 contracting cells and found a strong correlation between Vinc/E-cad ratio and cell area (Fig. 4K).

### Mechanism of Ca2+ induced cell contraction

Although we have observed that medio-apical myosin II accumulates in response to uncaging and that it correlates with the degree of cell contraction (Fig. 4A-E), the mechanism of how Ca^2+^ induces contraction is unclear. At least two different mechanisms are conceivable. First, Ca^2+^ may activate myosin II activity via the generic pathway involving Rho kinase and phosphorylation of the regulatory light chain. Secondly, Ca^2+^ may activate the myosin light chain kinase or directly engage at the acto-myosin filaments. We first tested whether the Ca^2+^ induced contraction depended on Rho kinase by employing its specific inhibitor Y-27632 [31]. We did not detect any cell contraction in embryos treated with the Rho kinase inhibitor indicating that Ca^2+^ induced contraction depends on Rho kinase (Fig. 5E and 5H, Supp. movie 7). Ca^2+^ uncaging was functional in these embryos (Fig. 5C and 5D, Supp. movie 6) as Ca^2+^ fluorescence in Y-27632 treated embryos was comparable in timing and magnitude to wild type embryos (Fig. 5D). The dependence on Rho kinase strongly supports the model that the Ca^2+^ signal acts via myosin II activation. These experiments rule out that Ca^2+^ induces a volume change independent of myosin II.

**Figure 5.**
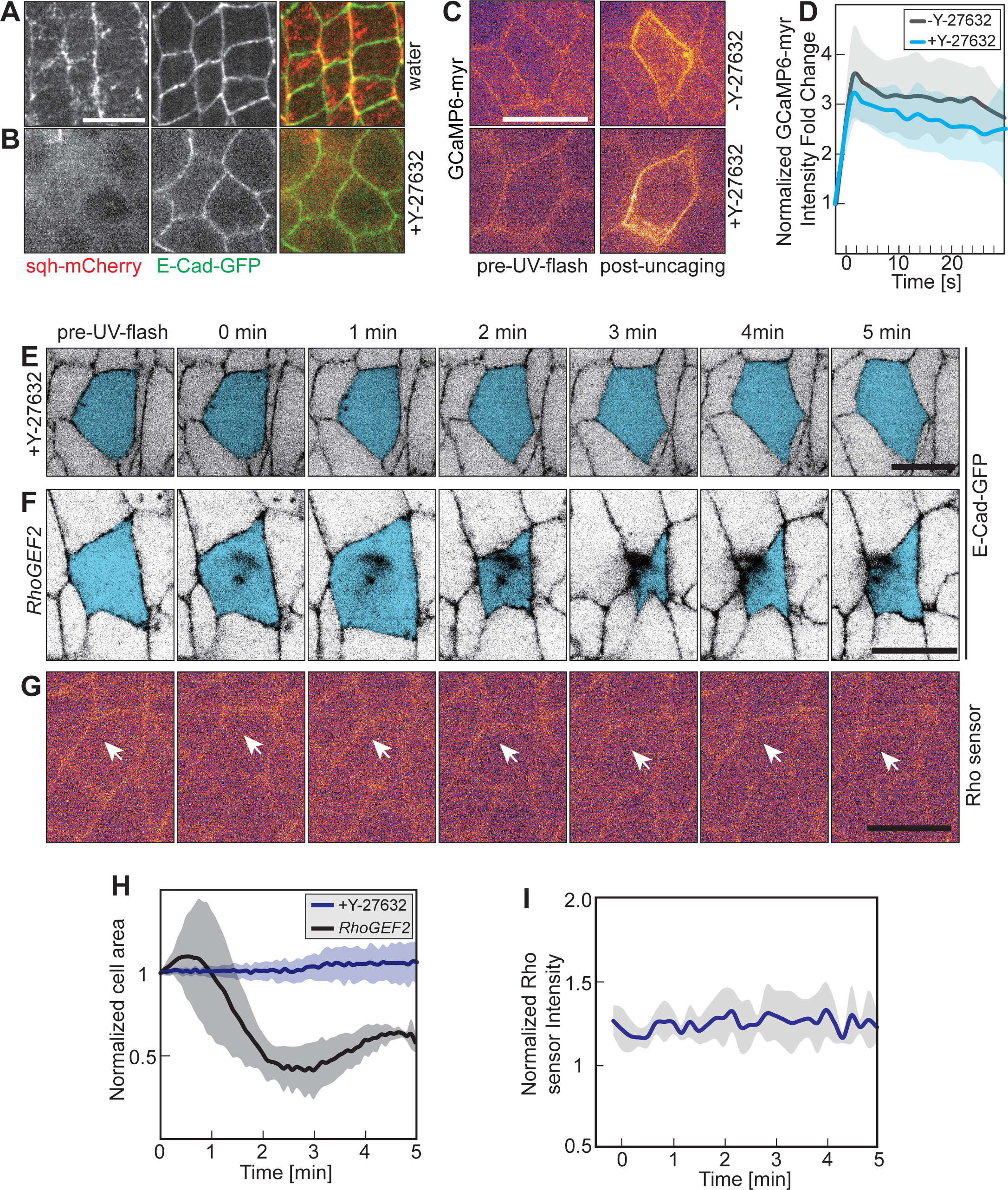
**Ca**^**2+**^ **induced constriction requires ROCK but not RhoGEF2.** **(A, B)** Images of embryos expressing sqh-mCherry and E-Cadherin-GFP injected with Y-27632 (ROCK inhibitor, 10 mM) or water. **(C, D)** Ca^2+^ uncaging in embryos (stage 7, lateral epidermis) expressing a membrane bound Ca2+ sensor (GCaMP6-myr) and injected with NP-EGTA,AM and Y-27632 as indicated. Images from time lapse recording. **(D)** Fluorescence intensity of GCaMP-myr in the target cell with (black, the same data with Fig. 1E) or without (blue) Y-27632. Mean (bold line) with standard deviation of the mean (ribbon band, 4 cells in 4 embryos). **(E–G)** Images from time-lapse recordings following Ca^2+^ uncaging (Lateral epidermis, stage 7). Taget cell marked in blue or by arrow in white. **(E)** Co-injected of Y-27632. **(F)** Embryos from *RhoGEF2* germline clones. **(G)** Embryo expressing a Rho sensor. **(H)** Cross sectional area of target cells normalized to initial size (the first frame of recording after uncaging) in embryos from *RhoGEF2* germline clones (black, n=4) and embryos injected with Y-27632 (blue, n=8) following Ca^2+^ uncaging. Mean (bold line) with standard deviation of the mean (ribbon band). **(I)** Rho sensor fluorescence in target cells in (n= 6) following Ca2+ uncaging. Scale bars 10 μm.

Rho kinase is activated by Rho signaling. RhoGEF2 is a major activator of Rho1 in the epidermal tissue during gastrulation, for example. We tested the dependence of the Ca^2+^ induced cell contraction on RhoGEF2 by conducting the uncaging in embryos lacking RhoGEF2. Quantification of the area dynamics of target cells revealed a behavior comparable in magnitude and timing to that in wild type embryos (Fig. 5F and 5H, Supp. movie 8). Lastly, we tested whether Rho1 was involved in mediating the Ca^2+^ signal to Rho kinase by visualizing Rho1 activation with a sensor protein. The Rho sensor was functional, since we detected an activation in cells undergoing cellularisation (Fig. S5A) and cytokinesis (Fig. S5B). In contrast, we did not detect a change in Rho sensor fluorescence in response to Ca^2+^ uncaging (Fig. 5G and 5I, Supp. movie 9). In summary, we propose a mechanism linking Ca^2+^ with myosin activation via Rho kinase but independent of Rho signaling via RhoGEF2.

## Discussion

We developed and validated Ca^2+^ uncaging as an optochemical method to induce cell contraction in epithelial tissues with precise temporal and spatial control. This approach enabled us to induce Ca^2+^ bursts in selected single cells within seconds inducing contraction to about half of the cross sectional area within a minute. The induced contraction did not damage cells or perturb tissue integrity. To our best knowledge, this is the first report for optically controlled cell contraction on the minute scale and at single cell resolution *in vivo* during epithelial tissue morphogenesis.

The optochemical system that we use is based UV laser photolysis of a photolabile Ca^2+^ chelator. The cage compound “NP-EGTA, AM” is membrane-permeant and thus allows convenient application to whole tissues. The 355nm UV laser we used is compatible with modern objectives and can be conveniently mounted on conventional live imaging microscopes. The dose of UV light depends on factors like light scattering by the tissue and thickness of the sample and needs to be carefully adjusted for the experimental system. Using a fluorescent Ca^2+^ sensor protein provides a direct optical readout of Ca^2+^ release and greatly aids UV intensity adjustment.

Importantly, Ca^2+^ uncaging does not require any genetically encoded protein and can be readily applied to any stock and genetic background. The independence from genetic constitution should vastly accelerate analysis and enable screening of mechanobiological cellular pathways and components, e. g. by comparing wide arrays of mutants to wild type behavior. In addition, Ca^2+^ uncaging is likely to open applications in manifold experimental systems with low genetic tractability. Importantly, UV Ca^2+^ uncaging leaves the entire visible spectrum available for optical interfacing with florescent protein indicators and opsin-based effectors. This in particular increases the options for simultaneously recording of cell and tissue behavior with the large palette of available fluorescent protein tags from CFP to RFP.

Ca^2+^ uncaging appears particularly suitable for studies of tissue morphogenesis. Specifically, intercellular coupling between neighboring cells poses a challenge to experimental design in studies of tissue morphogenesis. Here cause and consequence cannot be easily distinguished without targeted activation of cellular contractility and precise external control of cellular behaviors. Thus acute interference is mandatory for dissecting causal functional dependencies. Although the mechanistic details for Ca^2+^ induced contraction need to be further resolved, UV induced Ca^2+^ uncaging allows interference with the mechanics of tissues with appropriate temporal and high spatial resolution. The method can be applied to a wide range of processes and organisms and should greatly improve our ability to study the causal role of cell contractility and of tissue mechanics and mechanotransduction *in vivo*.

## Materials and Methods

### *Drosophila* strains and genetics

Fly stocks were obtained from the Bloomington Drosophila Stock Center, if not otherwise noted and genetic markers and annotations are described in Flybase [32]. Following transgenes were used UAS-GCaMP6-myr [20], E-Cadherin-GFP [33], E-Cadherin-mTomato [33],ubiquitin-E-Cadherin-GFP, Sqh-mCherry [25,34], Mat-Gal4-67,15 (D. St. Johnston, Cambridge/UK) and Amnioserosa-Gal4 (Bloomington), Rho sensor (R. Lehmann, New York/USA)

The allele *RhoGEF2[04291]* together with FRT[2R, G13] was recombined with ubi-E-Cad-GFP. *RhoGEF2* germline clones were generated and selected with ovo[D]. First and second instar larvae were heatshocked twice for 60 min at 37 °C.

### Cloning

Vinculin D1 domain (aa6–aa257) (HindIII-Xho1), eGFP (EcoR1-Xho1) were inserted between the EcoR1-Xho1 sites of a pUASt with attB sequence. PCR cloning was verified by sequencing of the fragments. pUASt-attB-VinculinD1-eGFP was inserted in chromosome II and recombined with E-Cad-mTomato. To get the expression of VinculinD1-GFP in amnioserosa tissue, the flies were crossed with AS-Gal4.

### Embryo preparation and injections

Embryos were prepared as described before [35]. Briefly, Embryos (2–2.5 hours at 25 °C in Fig. 1, 2 4A-4E and 5, and 15–17 hours at 20 °C in Fig. 3 and 4F-4K) were collected and dechorionated with 50% hypochloride bleach for 90 s, dried in a desiccation chamber for ∼10 min, covered with halocarbon oil and injected dorsally into vitelline space in the dark at room temperature (∼22 °C). After injection, the embryos were incubated at room temperature in the dark about 10 min prior to uncaging.

NP-EGTA, AM (Invitrogen) was prepared in 1X injection solution (180 mM NaCl, 10 mM HEPES, 5 mM KCl, 1 mM MgCl_2_ [pH 7.2]) [11]. 2 mM NP-EGTA, AM was injected for Ca^2+^ uncaging in epidermal cells, and 1 mM NP-EGTA, AM was injected for Ca^2+^ uncaging in amnioserosa cells. To inhibit Rock activity, 10 mM Y-27632 (Sigma) was injected.

### Ca^2+^ uncaging and imaging

We employed a 355 nm pulsed YAG laser (DPSL-355/14, Rapp Opto Electonic) mounted on the epiport. We illuminated under the ‘Click and Fire’ Mode on the ‘REO-SysCon-Zen’ platform (Rapp OptoElectonic), while a movie was recorded on a spinning disc microscope (Zeiss, 100x/oil, NA1.4) with a CCD camera. For the images in Fig. 2 and 4, the movies were recorded with an emCCD camera (Photometrics, Evolve 512) and the recording started about 20 seconds after Ca^2+^ uncaging. The intensity of the UV laser was adjusted so that no morphological changes were induced in 1X injection solution injected embryos. The laser was applied for 1.5 seconds (around 300 pulses) per cell with 2.5% laser power (∼0.5 mJ/cell).

To get the expression of GCaMP6-myr in the embryos, the flies were crossed with Mat-Gal4-67,15. The cross-section images were recorded in GFP channel with a frame rate of 1/s. For the images in Fig. 2, after uncaging, axial stacks of 3–4 images with 0.5 μm step size were recording in GFP channel with frame rate of 0.2/s (Fig. 2B-2E) or 0.1/s (Fig. 2F-2G). The recording started about 20 seconds after Ca^2+^ uncaging. For the images in Fig. 3 and 5E-G, the cross-section images were recorded in GFP channel with a frame rate of 0.2/s. To analyse myosin dynamics after Ca^2+^ uncaging (Fig. 4A and B), GFP and mCherry channels were recorded simultaneously with a frame rate of 0.1/s for E-Cad-GFP and Sqh-mCherry. After uncaging, axial stacks of 3–4 image with 1 μm step size were recording. The recording started about 20 seconds after Ca^2+^ uncaging. In Fig. 4G and 4F, GFP and mTomato channels were recorded simultaneously with a frame rate of 0.1/s. The apical plane of the embryo with 4 z-sections of 0.5 μm were acquired. In Fig. 5A and 5B, embryos expressing sqh-mCherry and E-Cad-GFP were injected with water or 10 mM Y-27632, GFP and mCherry channels were recorded simultaneously on a spinning disc microscope (Zeiss, 100x/oil, NA1.4) with an emCCD camera (Photometrics, Evolve 512). The apical plane of the embryo with 4 z-sections of 0.5 μm were acquired.

### Image processing and analysis

The fluorescence intensity of GCaMP6-myr (in Fig. 1 and Fig. 5) was measured manually with ImageJ/Fiji [36]. The integrated density (a.u.) was measured along the cell membrane and divided by the cell membrane length (μm) to get the mean fluorescence intensity *I*_*t*_. The background *I*_*b*_ was from the integrated density (a.u.) which are measured from the cytoplasm and divided by the measurement length (μm). The normalized GCaMP6-myr intensity fold increase was calculated as follows:

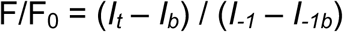

Where *I*_*t*_ is the mean intensity at time *t, I*_*b*_ is the mean intensity of the background at time *t*. *I*_*-1*_ is the mean intensity at 1 second before UV illumination, *I*_*-1b*_ is the mean intensity of the background at 1 second before UV illumination.

To analyse cell dynamics after Ca^2+^ uncaging, image stacks were projected by the “Max Intensity” option. The projected and cross-section images were segmented and tracked with “Tissue Analyzer” [37] in ImageJ/Fiji. Cell area measurements were carried out with ImageJ/Fiji. In movie 3, the Z-projected images were stabilized with “Image Stabilizer” [38].

To analyse myosin dynamics after Ca^2+^ uncaging (in Fig. 4), the image stacks from sqh-mCherry embryos were projected with the “Max Intensity” option. Mean medio-apical Sqh-mCherry fluorescence intensity was measured manually with ImageJ and normalized as follows:

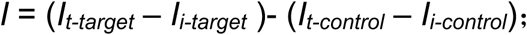

Where *I*_*t-target*_ is the fluorescence intensity of target cell at time t; *I*_*i-target*_ is the fluorescence intensity of target cell at the first frame after UV-illumination; *I*_*t-control*_ is the fluorescence intensity of control cell; *I*_*i-control*_ is the fluorescence intensity of control cell at the first frame after UV-illumination.

To analyse Rho sensor dynamics, the fluorescence intensity of Rho sensor (in Fig. 5I and S4) was measured manually with ImageJ/Fiji. The integrated intensity (a.u.) was measured along the cell membrane and divided by the cell membrane length (μm) to get the mean fluorescence intensity *I*_*t*_ (Fig. S5). The background *I*_*b*_ represents the averaged fluorescence intensity (a.u.) within the cytoplasm. The normalized Rho sensor intensity (in Fig. 5I) was calculated as follows:

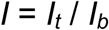

To analyse Vinculin D1 and E-Cadherin dynamics, the fluorescence intensity of VinculinD1-GFP (in Fig. 4I and S3) at cell junctions was measured manually with ImageJ/Fiji. The fluorescence intensity was measured along cell junctions and normalized to the initial fluorescence (t=0).

The ratio of VincuinD1-GFP/E-Cadherin-mTomato (in Fig. 4I and S3) was generated by plugin “Ratio plus” in ImageJ/Fiji. followed by the manual measurements of the intensity. The numbers represent the changes in relative fluorescence (Vinc-GFP/ECad-RFP) in reference to the initial fluorescence ratio (t=0):The Figures show the first 90 seconds (Fig 4I) and first 120 seconds (Fig S3).

## Acknowledgements

We are grateful to Marion Silies, Stefan Luschnig, Adam Martin, Ruth Lehmann and Daniel St Johnston for materials. We acknowledge service support from the Bloomington Drosophila Stock Center (supported by NIH P40OD018537). This work was in part supported by the Göttingen Centre for Molecular Biology (funds for equipment repair) and the Deutsche Forschungsgemeinschaft (DFG, FOR1756 GR1945/6-1/2, SFB937/TP10 and equipment grant INST1525/16-1 FUGG).

## Author contributions

DK conducted the experiments and analyzed the data. ZL generated the Vinculin-D1-GFP transgenic fly and analyzed the Vinculin-D1-GFP data. MH analyzed data and obtained in Fig. 2E 3F and 4E. FW and JG conceived and supervised the study. DK, FW and JG wrote the manuscript.

## Figure legends

**Figure S1 (related to Fig.2) Area traces of target cells.** Cross sectional cell areas of individual target cells (dashed lines) following a UV laser pulse. **(A)** Embryos injected with 2 mM NP-EGTA,AM (n=8 cells in 8 embryos). **(B)** Embryos injected with buffer (n=5 cells in 5 embryos). Mean (bold line) with standard deviation of the mean (ribbon).

**Figure S2 (related to Fig.4) VinculinD1 reporter.** Scheme of the domain structure of Vinculin. Numbers indicate position of amino acid residues. Transgenic construct with the D1 domain (blue) fused to GFP and expressed under GAL4/UAS control.

**Figure S3 (related to Fig.4) VinculinD1-GFP reporter in target cells. (A)** VinculinD1-GFP fluorescence on cell junctions of target and control cells (n=3 target cells and 3 control cell borders) **(B)** Scatter plot of normalized cross sectional area with normalized VinculinD1-GFP intensity in target cells.

**Figure S4 (related to Fig.5) The Rock inhibitor inhibits Ca**^**2+**^ **induced constriction.** Cross sectional areas of 8 individual target cells following Ca^2+^ uncaging in Y-27632 co-injected embryos.

**Figure S5 (related to Fig.5) Rho sensor is not increased following Ca**^**2+**^ **uncaging. (A)** Axial image stack of an embryo during cellularization showing the functionality of the Rho sensor. **(B)** Images from time lapse recording of an embryos expressing the Rho sensor. Cells in cytokinesins. Stage 8. Lateral epidermis. Scale bar 10 μm.

**Movie 1 Uncaging induces rapid intracellular Ca**^**2+**^ **concentration increasing in epithelial target cells.** Time-lapse recording from embryos expressing UAS-myr-GCaMP6 (lateral epidermis, stage 7). Time in minutes:seconds. Anterior left, dorsal up. This movie relates to Fig. 1C 1F and 5C.

**Movie 2 Ca**^**2+**^ **uncaging induces constriction in a columnar epithelium (including the process of uncaging, using CCD camera).** Time-lapse recording from embryos expressing E-Cad-GFP (lateral epidermis, stage 7) recorded with a CCD. The target cell is marked by a red dot. A stack was acquired every 5 seconds. Time is indicated as minutes: seconds. Anterior left, dorsal up.

**Movie 3 Ca**^**2+**^ **uncaging induces constriction in a columnar epithelium (without the process of uncaging, using emCCD camera).** Time-lapse recording from embryos were expressing E-Cad-GFP recored with a emCCD. The target cell is marked by a red dot. Stacks were acquired every 5 seconds. Time is indicated as minutes:seconds. Anterior left, dorsal up. This movie relates to Fig. 2C.

**Movie 4 Ca**^**2+**^ **uncaging triggers apical-constriction in a squamous epithelium.** Time-lapse recording of embryos expressing E-Cad-GFP (amnioserosa, stage 14). Stacks were acquired every 5 seconds. Time is indicated as minutes: seconds. Anterior left. This movie relates to Fig. 3D.

**Movie 5 VinculinD1-GFP accumulation following Ca**^**2+**^ **uncaging.** Time-lapse recording from amnioserosa cells in stage 14 embryos were expressing E-Cad-mTom and VinculinD1-GFP. Time is indicated as minutes:seconds. Anterior left. This movie relates to Fig. 4G and 4H.

**Movie 6 Y-27632 does not disturb Ca**^**2+**^ **uncaging.** Time lapse recording from embryos expressing UAS-myr-GCaMP6 and injected with 10 mM Y-27632 and 2 mM NP-EGTA,AM. A stack was acquired every second. Time is indicated as minutes:seconds. Anterior left, dorsal up. This movie relates to Fig. 5C.

**Movie 7 Ca**^**2+**^ **uncaging fails to trigger apical-constriction in Y-27632 injected embryos.** Time lapse recording from embryos were expressing E-Cad-GFP and co-injected with 10 mM Y-27632 and 2 mM NP-EGTA,AM (lateral epidermis, stage 7). The target cell is marked by a red dot. Time is indicated as minutes:seconds. Anterior left, dorsal up. This movie relates to Fig. 5E.

**Movie 8 *RhoGEF2* is not required for Ca**^**2+**^ **induced constriction.** Time lapse recording from embryos with RhoGEF2 germline clones expressing E-Cad-GFP (lateral epidermis, stage 7). Time is indicated as minutes: seconds. Anterior left, dorsal up. This movie relates to Fig. 5F.

**Movie 9 Rho A sensor activity is not induced by Ca**^**2+**^ **uncaging.** Time-lapse recording from embryos were expressing a Rho sensor (lateral epidermis, stage 7). The target cell is marked by a blue dot. Time is indicated as minutes: seconds. Anterior left, dorsal up. This movie relates to Fig. 5G.

