## Supplementary figures and images for "*In vivo* optochemical control of cell contractility at single cell resolution by Ca^2+^-mediated light-activation of myosin (CaLM)"

### Figure 2

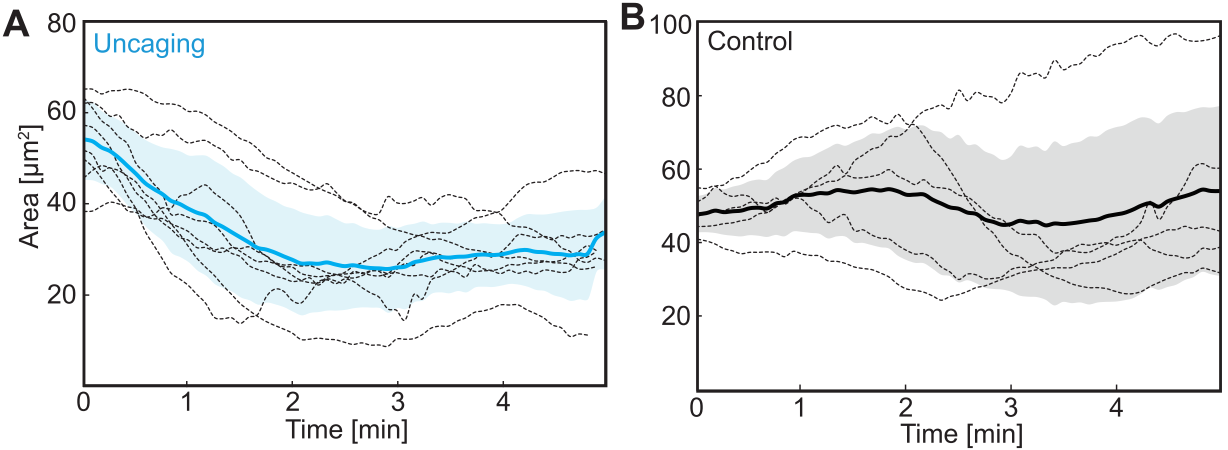

### Figure 4

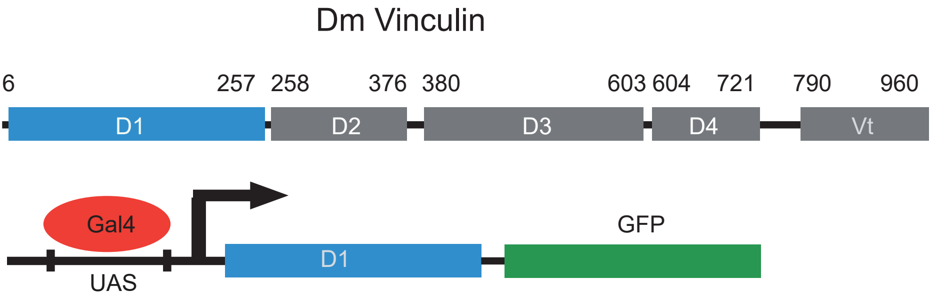

### Figure 4

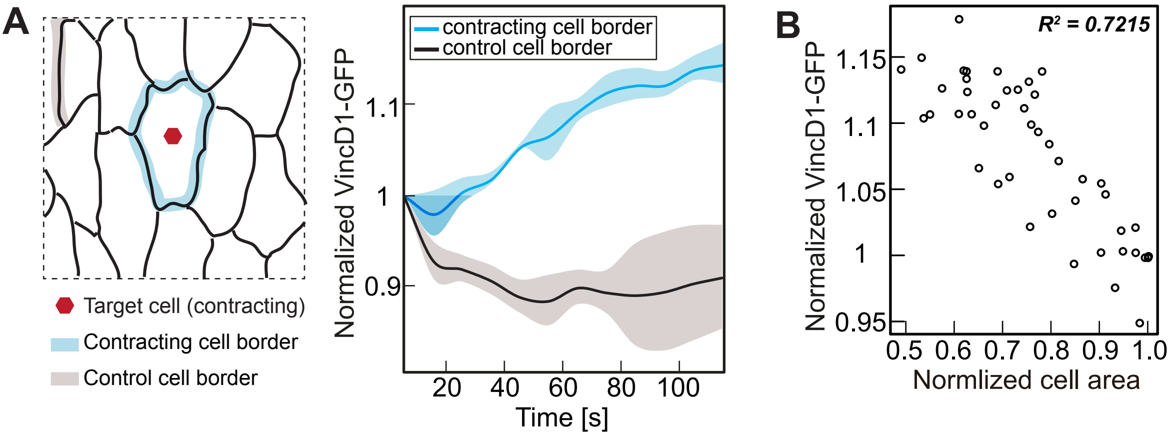
